# LongTron: Automated Analysis of Long Read Spliced Alignment Accuracy

**DOI:** 10.1101/2020.11.10.376871

**Authors:** Christopher Wilks, Michael C. Schatz

## Abstract

**Motivation:** Long read sequencing has increased the accuracy and completeness of assemblies of various organisms’ genomes in recent months. Similarly, spliced alignments of long read RNA sequencing hold the promise of delivering much longer transcripts of existing and novel isoforms in known genes without the need for error-prone transcript assemblies from short reads. However, low coverage and high-error rates potentially hamper the widespread adoption of long-read spliced alignments in annotation updates and isoform-level expression quantifications.

**Results:** Addressing these issues, we first develop a simulation of error modes for both Oxford Nanopore and PacBio CCS spliced-alignments. Based on this we train a Random Forest classifier to assign new long-read alignments to one of two error categories, a novel category, or label them as non-error. We use this classifier to label reads from the spliced-alignments of the popular aligner minimap2, run on three long read sequencing datasets, including NA12878 from Oxford Nanopore and PacBio CCS, as well as a PacBio SKBR3 cancer cell line. Finally, we compare the intron chains of the three long read alignments against individual splice sites, short read assemblies, and the output from the FLAIR pipeline on the same samples.

Our results demonstrate a substantial lack of precision in determining exact splice sites for long reads during alignment on both platforms while showing some benefit from postprocessing. This work motivates the need for both better aligners and additional post-alignment processing to adjust incorrectly called putative splice-sites and clarify novel transcripts support.

**Availability and implementation:** Source code for the random forest implemented in python is available at https://github.com/schatzlab/LongTron under the MIT license. The modified version of GffCompare used to construct Table 3 and related is here: https://github.com/ChristopherWilks/gffcompare/releases/tag/0.11.2LT

**Supplementary Information:** Supplementary notes and figures are available online.

## 1. Introduction and Background

The sequencing of cDNA derived from RNA molecules via fragmented reads, typically 75-250 base pairs long, known as short-read sequencing, has been extensively used for research for over 10 years (van Dijk et al., 2014). The RNA-seq approach has been utilized for determining gene expression (Bray et al., 2016; Patro et al., 2017), alternative gene structure (Dobin et al., 2013; Goldstein et al., 2016), and fusion constructs (Haas et al., 2019), as well as de novo regions of expression throughout genomes of different species (Trapnell et al., 2010; Pertea et al., 2015). Due to continued investment in improving these sequencers over time, the error rates of short reads are relatively low (<1%) while the throughput is high, with up to a terabase of data from a sequencer in less than 2 days.

Long read RNA sequencing is comparatively recent and has the capacity to complement or even surpass short read RNA-seq in its ability to span several exons and splice junctions of an isoform. This capability can in principle aid the discovery of novel isoforms and the expression of existing isoforms in specific tissues and cell types (**Figure 1**).

**Figure 1.**
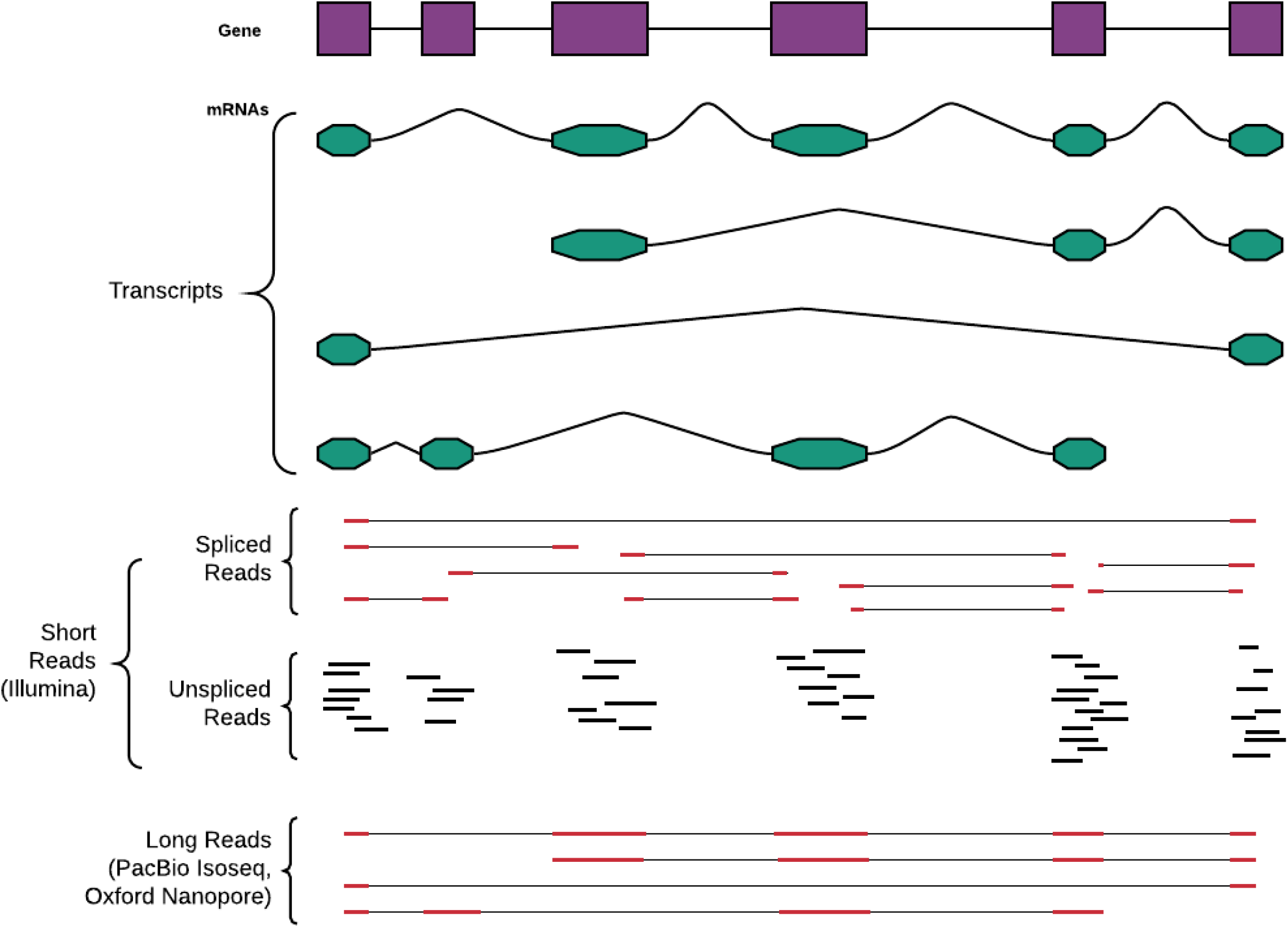
Long-reads versus short-reads. While short reads have much lower error rates (∼1% vs. ∼10%) and higher coverage they lack the general ability to connect multiple splicing interactions across the transcript due to their extreme shortness (250 bases vs. 10K’s bases).

However, long reads have a few problems currently including higher error rate, higher cost per read, and potential 3-prime end bias (Mantere et al., 2019; Amarasinghe et al., 2020). A fundamental challenge is that long reads suffer from a much higher error rate (2-10%) that is less systematic than the lower error rate of Illumina short reads (<1%, mostly toward the 3’ end of the read). Additionally, the likely lower throughput due to higher cost per read of long reads, may make transcript quantification and differential analyses more difficult across the transcriptome due to lower coverage at any given locus if whole transcriptome analyses are desired (Kovaka et al., 2019).

These problems have negative effects on downstream efforts aimed at understanding transcription, including the first step in most analyses, alignment. Recently, the multi-mode aligner, minimap2 (Li, 2018) was released and is gaining popularity in long read related work, both in DNA and RNA contexts. Minimap2 is fast and relatively accurate and these authors support its continued use. However, no aligner is perfect, and minimap2 does make mistakes, specifically in the areas of spliced-alignment and the mapping of long reads’ ends.

Thus this work is an initial attempt at studying and elucidating the cases where alignments of spliced long reads, both from PacBio IsoSeq and Oxford Nanopore DirectRNA, break down.The rest of the paper is divided into sections covering (1) the simulation of long reads and their alignments for benchmarking, (2) the random forest approach we took to predicting error categories of aligned long reads on both simulated and real datasets, and (3) the concordance between long reads and short reads with respect to individual splice junctions and whole transcripts, and (4) the results of our prediction approach run on real datasets.

### Related Work

This work extends the Qtip algorithm (Langmead, 2017) that also attempted to profile alignment quality/errors using a Random Forest. Where LongTron primarily differs is that we focus on the spliced alignment of long reads using minimap2 whereas Qtip focused on unspliced alignment of DNA short reads using Bowtie2 (Langmead and Salzberg, 2012), BWA-mem (Li, 2013), and SNAP (Zaharia *et al*., 2011) aligners.

Another related work is the FLAIR pipeline (Tang *et al*., 2018) which seeks to improve the spliced alignment of long reads. We utilized the FLAIR pipeline in our comparisons with raw minimap2 alignments in the results section of this paper. FLAIR uses known splice junctions from annotation and short read sequencing to correct and filter the set of spliced alignments for long reads. While FLAIR is a useful tool for correcting and refining the set of alignments, its use of annotated splice junctions makes it potentially problematic for studies looking for novel splicing in long read alignments. A related pipeline similarly profiling long reads, specifically for the PacBio platform is SQANTI (Tardaguila *et al*., 2018). SQUANTI and its successor SQUANTI2 (https://github.com/Magdoll/SQANTI2/) are intended to classify PacBio long reads spliced alignments and also use a Random Forest to classify artifactual results.

## 2. Methods

### 2.1 Long read failure modes

Typically RNA-seq aligners leverage heuristics to find a set of near-optimal candidate locations in the genome for the placement of both short and long reads. For RNA sequence analysis these heuristics are particularly relevant for at least two phases of the alignment process, commonly called seed-and-extend. In the first phase, the alignment search space is narrowed down from being the full genome to a short list of candidate loci (using seeds). These seeds are chosen in different ways by various aligners, although they often use heuristics that don’t guarantee an optimal alignment will always be identified (Darby et al., 2020). In the second phase, candidate loci are more thoroughly checked for their compatibility with the query sequence which includes splice-site determination. The Smith-Waterman optimal algorithm (Smith and Waterman, 1981) for local sequence alignment can be used efficiently at this stage to produce a gapped alignment. However, this is not useful for spliced alignments which still require a heuristic to determine the best splice-sites around which to split the query sequence.

### 2.2 Long Read Transcriptome Simulation

To assess the accuracy of a long read RNA-seq analysis pipeline, we first used a simulation approach so that we could precisely measure the alignment accuracy and splicing results of the simulated reads compared to their ground truth. For this, we started with the Gencode version 28 annotation and the error profile from SURVIVOR (Jeffares *et al*., 2017) for both Oxford Nanopore and PacBio IsoSeq derived from minimap2 alignments of NA12878 reads to the Gencode transcriptome. The NA12878 sample is from a disease-free human and has been used by many other research efforts. Using these we simulated long reads from the transcript sequences, both full-length and partial length. We then aligned these simulated reads against the genome and extracted features from each alignment. These features were then evaluated by a random forest for training and prediction. Our implementation used the *RandomForestClassifier* in the scikit-learn Python machine learning framework. We used 100 trees and eight parallel threads for training. For the purposes of Receiver Operator Curve (ROC) plotting we used the *predict_proba* method on the held-out test set. The genomic alignments of the simulated reads were used to determine four correctness categories. We experimented with using both these four categories as a multi-class prediction problem in the random forest as well as a more simple binary model where the three non problem-free categories in the list below were collapsed into one category.

Junction alignments were first categorized into five subcategories, allowing a margin (“fuzz”) of up to 20 nuclotides on each end, as described in **Figure 2.2**. These five categories were then categorized into the four top-level correctness classes:

**Figure 2.1.**
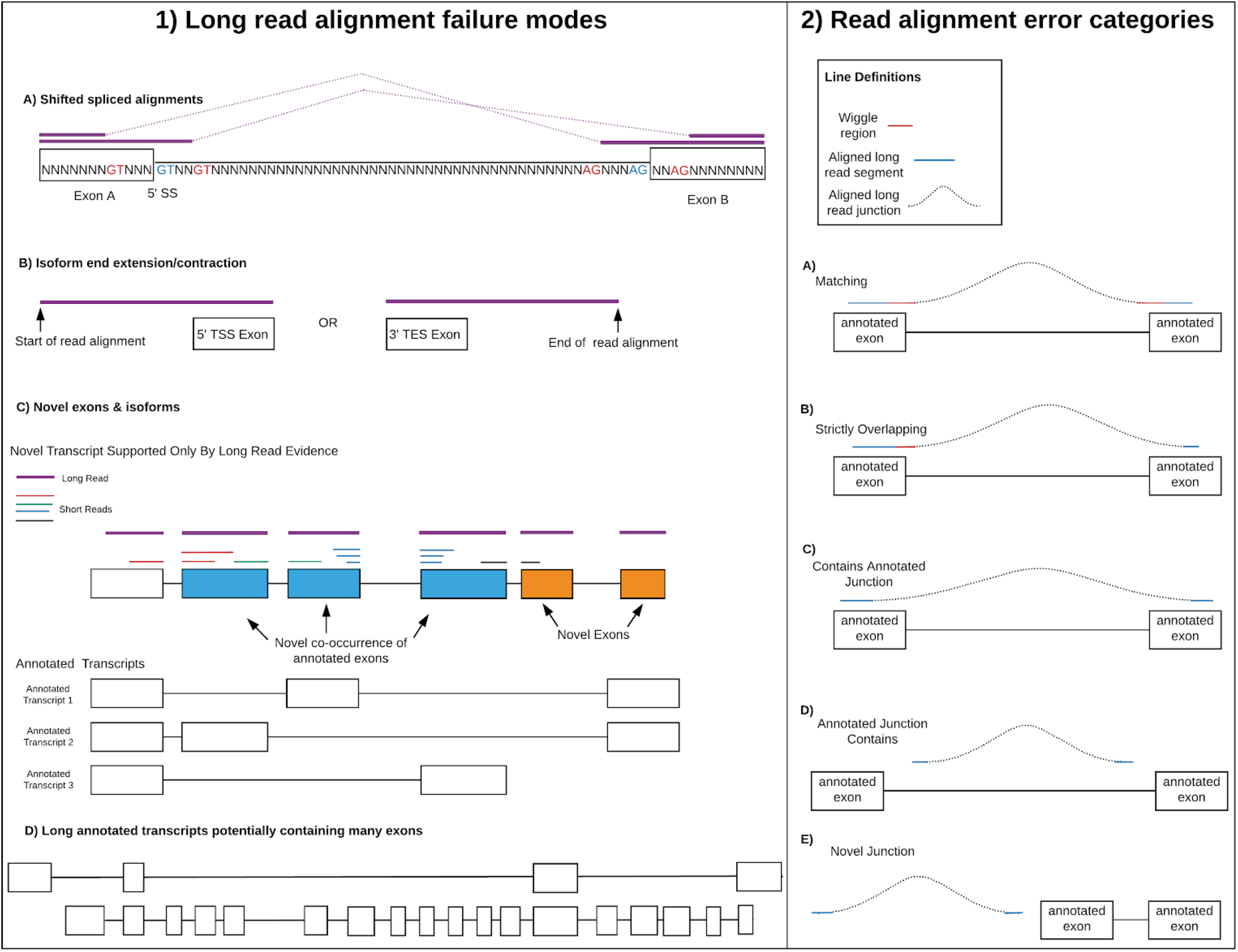
Long read alignment failure modes. A) Spliced alignments can shift in the presence of unannotated splice motifs in the reference near annotated (real) splice sites. B) 5’ and 3’ ends of isoforms are difficult to get right as sequencing the ends of long reads is imprecise. C) Long reads can produce novel configurations of annotated exons and/or novel exons. However, these may be simply alignment artifacts due to splice motifs and/or repeats in the region (e.g. the rightmost novel exon has no short read support). D) Large numbers of exons (splice sites) can result in multiple novel long read alignments, some of which may be false. This is in part due to the non-full length nature of many of the long reads (especially from PacBio). **2**.**2. Read alignment error categories**. A) Matching junction alignment against at least one source transcript junction; B) Alignment overlapping any transcripts’ junction; C) Alignment containing any transcripts’ junctions; D) One or more transcripts’ junctions containing aligned junction; E) Junction is completely novel

1. Problem-free (A)
2. Any error (alignment in any of B-D but not all three)
3. Recurrent error (alignments in all three B-D)
4. Novel (E)

With the exception of splice motifs as the third most important feature in the Oxford full-length run, exon length dominated the Oxford feature importance rankings (**Table S1**). Similarly, both exon and transcript length were among several of the top most important features for PacBio. In addition GC content was the third most important feature for the PacBio full-length run. A selection of these features are shown in **Figure 3.2**.

**Figure 3-1.**
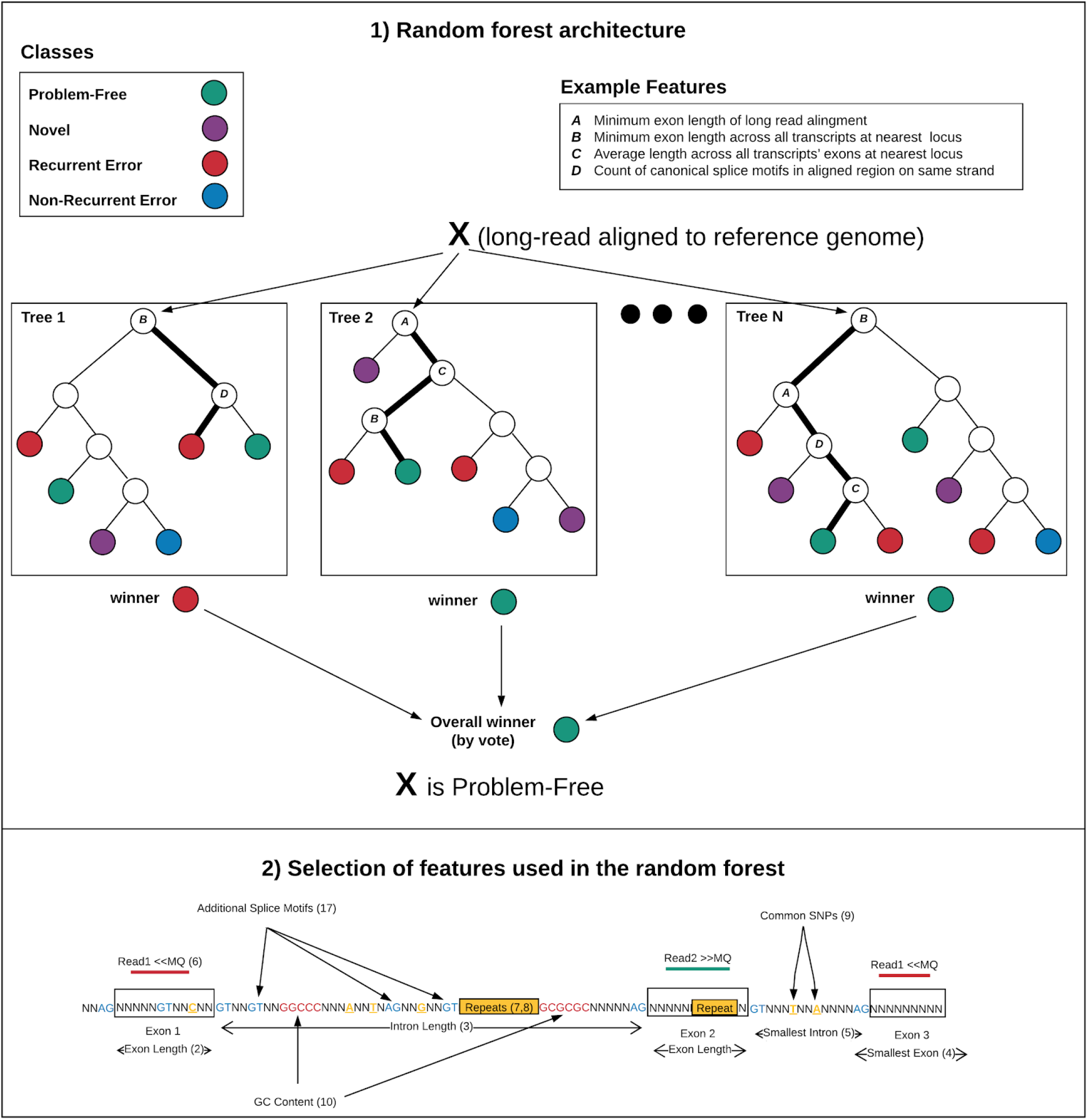
Random forest classification. **3-2**. Diagram of a selection of features used in the random forest, including 1-10 and 17 from Table S2.

## 3. Results

We trained four distinct random forest models using the final set of features described above:

1. PacBio IsoSeq Full Length
2. PacBio IsoSeq Fragment
3. Oxford Nanopore Full Length
4. Oxford Nanopore Fragment

Training accuracy was high on a held out test dataset (Supplement Figures **S2A-D**).

We then applied both full- and fragment-length models to the minimap2 alignments of long reads from PacBio and Oxford sequencing of the NA12878 sample. We intersected the long reads alignments with transcripts of known error categories to get the ground truth using BEDTools (Quinlan et al., 2010). This allows us to compute a form of recall and precision of the predictions (**Supplemental Tables S3 and S4**).

These results show Oxford had more errors than PacBio and also full-length alignments are more difficult to achieve than are fragments (**Table 1**). The PacBio IsoSeq platform supports the ability to generate a set of higher quality long reads by continuing to sequence the same molecule iteratively in a process called Circular Consensus Sequencing (CCS) (Gordon *et al*., 2015). The NA12878 PacBio dataset we are using is CCS corrected which likely contributes to its higher problem-free percentages.

**Table 1.** Counts of alignments in each simulated training class.

| Dataset | Total | Problem-free | Any error | Recurrent error | Novel |
| --- | --- | --- | --- | --- | --- |
| Oxford full | 1,696,509 | 41.90% | 55.42% | 2.30% | 0.38% (6,511) |
| **length** |  | (710,869) | (940,159) | (38,970) |  |
| **Oxford fragment** | 1,668,351 | 75.09%<br>(1,252,754) | 19.64%<br>(327,629) | 2.29%<br>(38,252) | 2.98% (49,716) |
| **PacBio CCS full length** | 971,786 | 88.37%<br>(858,755) | 9.75%<br>(94,708) | 1.33%<br>(12,883) | 0.56% (5,440) |
| **PacBio CCS fragment** | 956,945 | 91.98%<br>(880,189) | 6.57%<br>(62,837) | 0.72% (6,900) | 0.73% (7,019) |

### 3.2 Splice-junction and Isoform Comparison

A significant portion of the work described here involved comparison of splicing and isoforms across both long read sequencing approaches as well as Illumina short read sequencing. In **Table 2**, we present a comparison of splice junctions between the three long read sequencing samples we used and a large compendium of putative splice junctions called from Illumina short reads, used in the Snaptron tool (Wilks *et al*., 2017).

**Table 2.** Splice Junction Comparison (Snaptron represents a compendium of short-read derived junctions, annotated and novel), fuzz=20 for bases on either side, percents do not add up to 100 as annotated short-reads are a subset of all short-reads. Junctions are compared by coordinates alone (strand not included).

| Long Read Sample | Total | Gencode V29 exact<br>(360,700) | Annotated short-read supported exact | All short-read supported exact<br>(111,160,460) | Fully novel exact | Gencode V29 fuzz | All short-read supported fuzz | Fully novel fuzz |
| --- | --- | --- | --- | --- | --- | --- | --- | --- |
|  |  |  | (Snaptron,<br>445,651) |  |  |  |  |  |
| PacBio<br>NA12878 | 301,666<br>(100%) | 96,124<br>(32%) | 99,394<br>(33%) | 115,496<br>(38%) | 186,170<br>(62%) | 258,094<br>(86%) | 292,751<br>(97%) | 8,915<br>(3%) |
| Oxford<br>NA12878 | 612,614<br>(100%) | 163,199<br>(27%) | 180,028<br>(29%) | 338,773<br>(55%) | 273,841<br>(45%) | 306,045<br>(50%) | 508,798<br>(83%) | 103,816<br>(17%) |
| PacBio<br>SKBR3 | 1,639,588<br>(100%) | 143,371<br>(9%) | 152,346<br>(9%) | 242,903<br>(15%) | 1,396,685<br>(85%) | 1,001,378<br>(61%) | 1,271,853<br>(78%) | 367,735<br>(22%) |
| Gencode<br>V29 | 360,700<br>(100%) | NA | NA | 346,810<br>(96%) | NA | NA | 350,671<br>(97%) | 10,029<br>(3%) |

As seen in the table requiring exact matches between the aligned long reads and the short-read based splice sites results in a minority of splice junctions matching in the two annotated categories. Allowing for a “fuzz” (20 bp) on both ends of a match greatly increases concordance between long reads and short reads. This discrepancy between exact and fuzz matching, specifically with annotated junctions, highlights the difficulties in long read alignment discussed in this paper.

In contrast, the “All short-read supported exact” and “Full novel exact” categories fare considerably better in concordance. This is one benefit of using a large group of short-read-derived splice junctions which include many putative novel junctions. Most of these matches would have been missed if a pseudo-alignment/quasi-mapping strategy had been used to derive the short read junctions.

Overall, the results from splice-junction concordance is relatively positive. While long read alignments fail to pick up many annotated junctions under the strictures of exact matching, the majority of them are relatively close in terms of genomic coordinates. Further, there is evidence here to suggest long read alignments are supporting other, novel junctions previously found in short reads.

Another area of importance is isoform level concordance. We’ve already noted the potential benefit of long reads in finding novel isoforms where these can either use novel exons/splice junctions, or more likely, novel groupings of existing exons/splice junctions. **Table 3** presents the isoform-level comparison results using intron-chains as a proxy for isoforms. Intron chains, as their name implies, restrict comparison to the order and identity of the genomic coordinates which make up the donor/acceptor sites within the isoform. Thus start/end coordinates of the isoform as a whole are ignored. This will miss differences arising from alternative transcript start/end sites although these are intrinsically the most difficult to sequence because of the protocols involved (Workman *et al*., 2019; Roach *et al*., 2020).

**Table 3.** Isoform comparison table, using gene models from Gencode V29, plus the isoforms from all the union of annotations; both exact and fuzz comparisons of the set of long-read derived isoforms which 1) match in number of introns or 2) are contained or contain a reference isoform. Percentages use the “Total Intron Chains” as the denominator for the row.

| Sample | Total<br>Intron<br>Chains | Gencode V29<br>(199381) | Union of<br>Annotations<br>(1098511) | Short Assembly | (Other) Long Reads |
| --- | --- | --- | --- | --- | --- |
| PacBio SKBR3<br>(PB-SKBR3) | 1,339,872 | 22.0%<br>(60.2%) | 27.7%<br>(63.0%) | Illumina: 16.7%<br>(44.2%) | NA |
| PacBio<br>NA12878 FLAIR | 13,026 | 87.1%<br>(95.6%) | 93.3%<br>(98.1%) | Illumina: 76.6%<br>(86.9%) | OX-FLAIR: 86.4% (92.3%),<br>PB-RAW: 93.5% (97.1%) |
| PacBio<br>NA12878<br>(PB-RAW) | 516,021 | 41.4%<br>(81.0%) | 43.4%<br>(76.9%) | Illumina: 39.0%<br>(64.6%) | OX-RAW: **61.4% (94.6%)**,<br>PB-FLAIR: 38.6% (75.1%) |
| **Oxford NA12878 FLAIR** | 43,817 | 65.7% (86.5%) | 79.8% (93.3%) | Illumina: 42.4% (62.5%) | OX-RAW: 98.5% (100.0%),<br>PB-FLAIR: 30.1% (42.7%) |
| **Oxford NA12878 (OX-RAW)** | 6,744,568 | 73.2% (89.1%) | 77.2% (90.9%) | Illumina: 67.4% (**83.5%**) | OX-FLAIR: 73.4% (87.8%),<br>**PB-RAW: 87.6% (94.5%)** |
| **Illumina SKBR3** | 30,866 | 63.2% (81.0%) | 77.6% (90.1%) | NA | PB-SKBR3: 69.9% (81.4%) |
| **Illumina NA12878** | 101,151 | 37.5% (57.6%) | 56.7% (73.5%) | NA | OX-RAW: 63.9% (**72.3%**),<br>OX-FLAIR: 21.9% (33.2%),<br>**PB-RAW: 39.0% (46.8%)**,<br>PB-FLAIR: 16.4% (21.4%) |

The totals column in **Table 3** represents deduplicated sets of intron chains (additional details in **Supplemental Note 2 and Table S5**). The intron-chains between samples show little concordance when exact matching is required. This phenomenon was initially worse than in the splice-junction level analysis (**Table 2**). These initial results spurred us to involve an additional approach to use in filtering, the FLAIR (Tang *et al*., 2018) pipeline. It also required us to modify an existing tool, gffcompare (Pertea et al., 2020), to allow for fuzz when comparing intron chains.

One issue we encountered was the ambiguity of strand of origin for the PacBio long reads. We noticed that a large number of mismatching PacBio read alignments were classified as matching but on the opposite strand when compared with the “Union of Annotations” transcript set. By considering the PacBio alignments which were classified by gffcompare as opposite strand matches (categories “o” and “s”) and swapping their strands, and then re-comparing, a larger number of alignments were correctly re-classified for PacBio. In contrast, changing the strand parameter (“-u”) in minimap2 had little effect. This is an important issue to consider when aligning PacBio-derived long reads with minimap2.

A key finding in **Table 3** is that allowing for fuzz around junction boundaries makes a substantial contribution to raising the number of matching intron chains across almost every category. This again underscores one of the key problems in long read alignment, that is any difficulties computing the exact coordinates of a single junction correctly are magnified when chaining together multiple of those junctions into isoforms.

However, even without fuzz, both Oxford (NA12878) and PacBio (SKBR3) aligned samples are able to capture a larger amount of the annotated intron chains than their short read assembled counterparts (Illumina NA12878/SKBR3). The fact that the PacBio NA12878 sample falls behind here may be due to the much lower numbers of reads present in that sample. This bodes well for long read sequencing in the future in terms of finding coverage for annotated isoforms. In addition, the FLAIR pipeline raises concordance dramatically but at the cost of a substantial reduction in total isoforms.

Further, even when requiring exact junction coordinate matches, the concordance between Oxford and PacBio is fairly high (61.4% and 87.6% respectively), while with a fuzz of 20bp the numbers both jump to ∼94% of each set. The lower percent of Oxford captured by PacBio is most likely due to the much smaller size of the PacBio read set. This is also probably the explanation for the lower percentage of Illumina-assembled intron chains captured by the NA12878 PacBio set (46.8%) even with fuzz, while the short read assembly is capturing a majority of the PacBio long read intron chains (64.6%) with fuzz. Oxford in comparison, is both being captured by and is capturing, at a high rate, the Illumina assemblies (83.5% and 72.3% with fuzz, respectively).

### 3.3 Effects of Random Forest Classifier on Transcript Matching against the Annotation

We further took the set of NA12878 (Pacbio and Nanopore) and SKBR3 alignments predicted to be in the “Problem-Free” category and used them in comparisons against the “Union of Annotation” transcript set to see if the number of matching intron-chains improved (**Table S6**). While this strategy improved the precision, the percent of total query read alignments which matched an annotated transcript (NA12878 or SKBR3), it substantially lowered the recall, the percent of annotated transcripts matching a query read alignment. This was in large part due to using the set of predictions where both the full-length and the fragment models predicted a problem-free alignment. Recomputing the comparison with the union of full-length and fragment models’ predictions results in close to the original recall while only slightly lowering the improved precision for Nanopore, but with less impact on PacBio.

### 3.4. Novel Alignment Examples in NA1878 and SKBR3

We next evaluated the use of long read RNA sequencing to discover novel (unannotated) transcripts in the genome (**Figure 4)**. In **Figure 4a**, both Oxford and PacBio long-reads from the NA12878 sample support some additional transcription before the start of the NPIPB5 gene using the UCSC Genome Browser (Haeussler et al., 2019). This could be a novel alternative transcription start site (TSS). The substantially reduced support from PacBio reads could be a factor due to the much smaller total read set in that sequencing experiment. **Figure 4b** displays a region of potential novel transcription found primarily in the SKBR3 PacBio long-read sample. While a small subset of the region has minor support in the Oxford sequenced NA12878 sample, the majority of the transcription appears to be exclusive to SKBR3. There appears to be further evidence from human mRNA/ESTs that this region is indeed transcribed and is not due to technical error

**Figure 4a.**
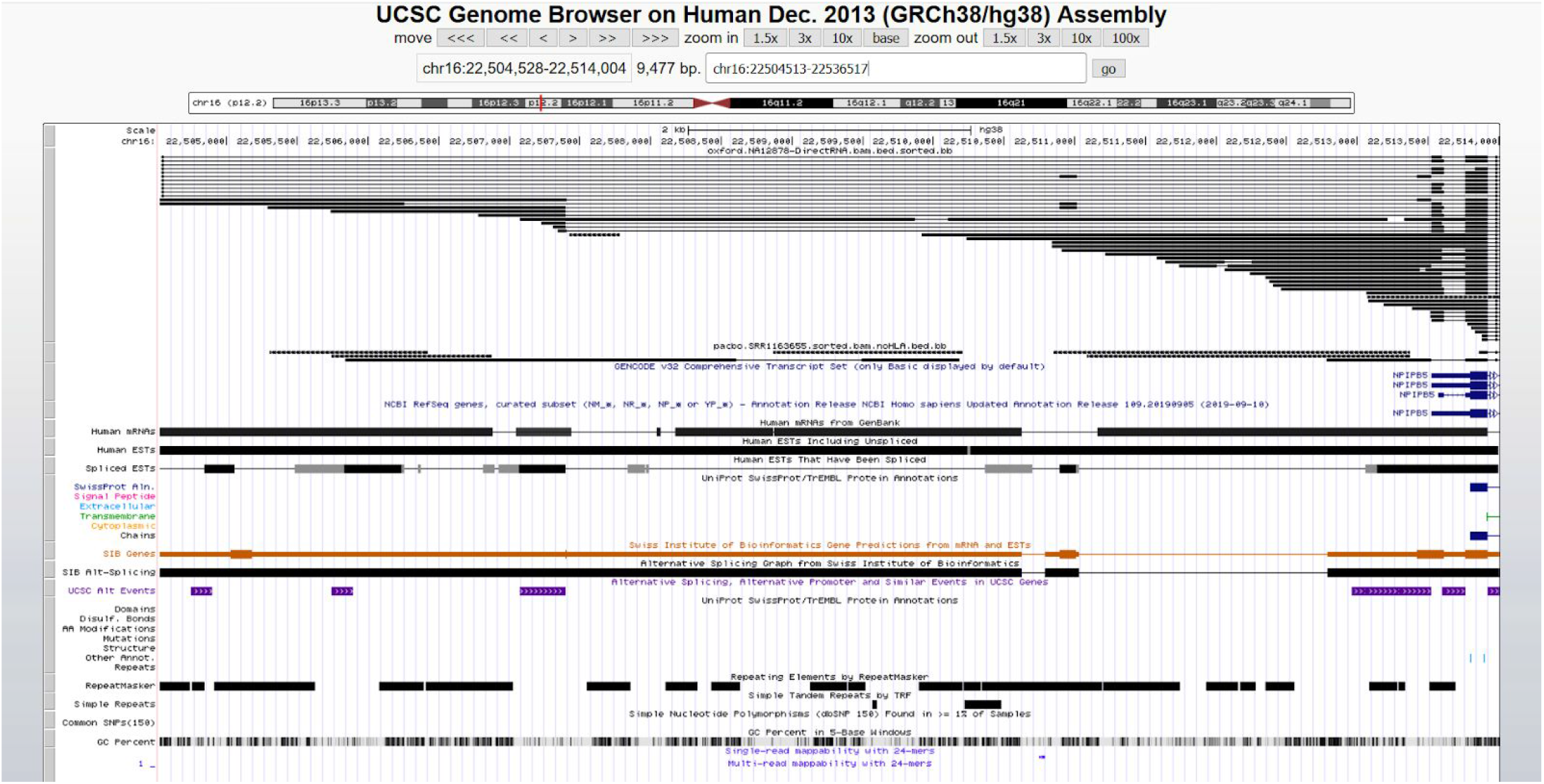
Novel transcript predicted region on NA12878 for both Oxford and PacBio

**Figure 4b.**
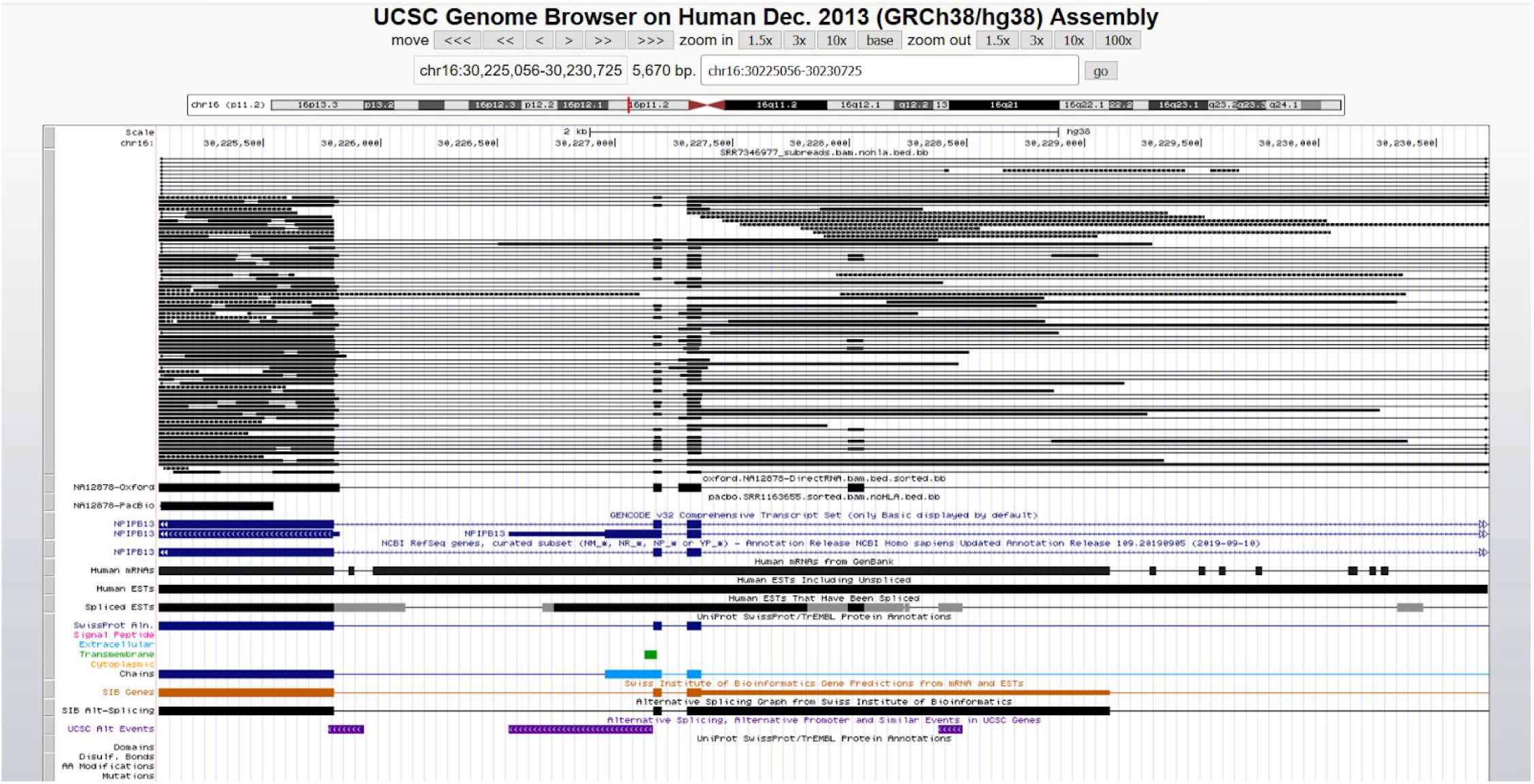
Novel transcript predicted region on SKBR3 PacBio

## 4. Discussion

Long reads are useful for finding new isoforms as combinations of splice junctions that have already been found by short reads, but caution must be exercised due to the failure modes described here. Our investigation will help in assessing long read alignments to make more confident calls as to 1) errors and 2) novel cases.

While there are a number of potential factors influencing how long reads are aligned, based on our investigation and the results of our random forest experiments, a few rise to the top in terms of importance. An important factor is the number of exons present in a gene. If there are more exons in an isoform, that translates into a larger number of potential splice-site determination errors the aligner can make when aligning long reads, which often are still fragments of the full length isoform. A related factor is the number of alternative isoforms present in the gene. This also raises the potential for splice-site finding errors in the aligner as many exons may be shared, while others may overlap but with different starts/ends, while still others are completely novel. This can lead to long reads missing certain alternative splice sites while supporting others within the same gene.

Further, the significant decrease in coverage within the currently available long-read sequencing datasets substantially reduces confidence in putative novel regions. This is partly alleviated by short reads, at least for single exon genes and combinations of a few splice junctions. However, long novel transcripts combining many exons in new ways will be harder to substantiate. It’s also important to consider that nucleotide sequence alignment in general is almost always heuristic-based, and this is certainly true of spliced-alignment. While better alignment heuristics, modeling, and short-read sequencing may be able to fill in some of the gaps left in long-read spliced-alignments, ultimately there will need to be either a significant decrease in error rates or a substantial increase in coverage to alleviate at least some of the problems reported here.

## Supporting information

LongTron Supplement

## Acknowledgements

The authors would like to thank Ben Langmead for his support of the project throughout and his insights. The authors would also like to thank Geo Pertea for his input on using his tool, GffCompare in this project. This research project was conducted using computational resources at the Maryland Advanced Research Computing Center (MARCC).

