## Supplementary material for "LongTron: Automated Analysis of Long Read Spliced Alignment Accuracy": LongTron Supplement

### Supplemental Materials

**Supplemental Table S1. Top 5 Most Important Features by Category.** 2

**Supplemental Table S2. Complete List of Random Forest Features** 4

**Supplemental Note 1. Details on junction matching** 6

**Supplemental Note 2. gffcompare run details** 7

**Supplemental Note 3. Training simulation dataset pipeline** 8

**Supplemental Note 4. Counting results of predictions on NA12878** 9

**Supplemental Note 5. NA12878 & SKBR Custom Tracks in the UCSC Genome Browser** 10

**Supplemental Note 6. Features used in the Random Forest training/prediction** 11

**Supplemental Table S1. Top 5 Most Important Features by Category.**

FL=full length, nFL= fragment

| Category | First | Second | Third | Fourth | Fifth |
|---|---|---|---|---|---|
| **Oxford FL 4 class** | Smallest aligned segment size (exon) (10%) | Minimum exon length across all transcripts at gene locus (6%) | Aggregate intron length (5%) | Count of canonical splice motifs in region on the same strand (4%) | Log of aggregate exon length (#2) (4%) |
| **Oxford FL 2 class** | Smallest aligned segment size (exon) (12%) | Minimum exon length across all transcripts at gene locus (7%) | Aggregate intron length (5%) | Count of canonical splice motifs in region on the same strand (4%) | Log of aggregate exon length (#2) (4%) |
| **Oxford nFL 4 class** | Minimum exon length across all transcripts at gene locus (10%) | Average length across all transcripts. exons at gene locus (6%) | Average ratio of non-unique k-mers per transcript base pair within transcripts which overlap target region (4%) | Maximum exon length across all transcripts at gene locus (4%) | Minimum transcript length across all transcripts at gene locus (4%) |
| **Oxford nFL 2 class** | Minimum exon length across all transcripts at gene locus (11%) | Average length across all transcripts. exons at gene locus (6%) | Maximum exon length across all transcripts at gene locus (4%) | Average ratio of non-unique k-mers per transcript base pair within transcripts which overlap target region (4%) | Minimum transcript length across all transcripts at gene locus (4%) |
| **PacBio FL 4 class** | Smallest aligned segment size (exon) (18%) | Minimum exon length across all transcripts at gene locus (9%) | Per-base average of GC content score in region (3%) | Aggregate intron length (3%) | Per-base average of overlapping exons (.transcript density.) (3%) |
| **PacBio FL 2 class** | Smallest aligned segment size (exon) (20%) | Minimum exon length across all transcripts at gene locus (9%) | Per-base average of GC content score in region (3%) | Aggregate intron length (3%) | Per-base average of overlapping exons (.transcript density.) (3%) |

| | | | | | |
|---|---|---|---|---|---|
| **PacBio nFL 4 class** | Minimum exon length across all transcripts at gene locus (11%) | Average length across all transcripts. exons at gene locus (5%) | Minimum transcript length across all transcripts at gene locus (5%) | Sum of all transcript lengths (transcript length=sum of exon lengths in transcript), this could be redundant across transcripts (5%) | Average ratio of non-unique k-mers per transcript base pair within transcripts which overlap target region (4%) |
| **PacBio nFL 2 class** | Minimum exon length across all transcripts at gene locus (12%) | Average length across all transcripts. exons at gene locus (6%) | Minimum transcript length across all transcripts at gene locus (5%) | Sum of all transcript lengths (transcript length=sum of exon lengths in transcript), this could be redundant across transcripts (5%) | Maximum exon length across all transcripts at gene locus (5%) |

**Supplemental Table S2. Complete List of Random Forest Features**

**General Features**

1. Sequence read length (including softclipping)
2. # of exons
3. Aggregate exon length
4. Aggregate intron length
5. Smallest aligned segment size (exon)
6. Smallest intron size
7. Mapping quality
8. # bases overlapping with RepeatMasker annotation
9. # bases overlapping with simple repeats (from Tandem Repeat Finder, subset of #7?)
10. Count of overlapping common 150 SNPs

**Means of region content**

11. Count of overlapping transcripts/reads on the same strand, always has at least 1 (itself)
12. Per-base average of GC content score in region
13. Per-base average of Multi-track Mappability score (k=24, umap) in region
14. Per-base average of overlapping exons ("exon density")
15. Per-base average of overlapping exons ("transcript density")

**Log of first 3 stats**

16. Log of # of exons (#1)
17. Log of aggregate exon length (#2)
18. Log of aggregate intron length (#3)

**Splice motifs & Segmental Duplications**

19. Count of canonical splice motifs in region on the same strand
20. Count of overlapping segmental duplicates
21. Ratio of the region that overlaps segmental duplicates (by base)

**Local mappability stats (k=10)**

22. Average of overlapping transcripts' base pair length
23. Average # of unique k-mers within transcripts which overlap target region
24. Average # of non-unique k-mers within transcripts which overlap target region
25. Average ratio of non-unique k-mers per transcript base pair within transcripts which overlap target region

**Closest Gene Locus Statistics**

26. # of transcripts at gene locus
27. Sum of all transcript lengths (transcript length=sum of exon lengths in transcript), this could be redundant across transcripts
28. Minimum transcript length across all transcripts at gene locus

29. Maximum transcript length across all transcripts at gene locus
30. Average transcript length across all transcript at gene locus
31. Total number of exons across all transcript at gene locus
32. Minimum exon length across all transcripts at gene locus
33. Maximum exon length across all transcripts at gene locus
34. Average length across all transcripts’ exons at gene locus
35. Distance to closest gene locus in base pairs (can be 0 or negative if closest locus is upstream)

**Supplemental Note 1. Details on junction matching**

**Novel:** read junctions have no overlap even within a fuzz of any annotated junction (no overlap either, so these aren't within any annotated junction). These are removed from consideration in the error categories below.

**Matching**:
Junction matching criteria:
1) Each end must be within the window of its annotated end
2)
a) at least one of the aligned jx's read IDs must match the annotated transcript ID it's overlapping
b) exon/jx idx must match within the overlapping transcript

The non-match from above are further split into categories:

**Overlapping:** a read junction which strictly overlaps an annotated junction (no containment for either).

**Contained (read junction)**: read's junction is either fully within an annotated junction OR its ends extend beyond the annotated junction's ends, but not beyond fuzz distance of the annotated junction's ends.

**Contained (annotated junction)**: read's junction ends are beyond the annotated junction's ends and both beyond fuzz distance of the annotated junction's ends.

**Supplemental Note 2. gffcompare run details**

In order to perform an accurate comparison at the isoform level we determined that we needed to modify an existing tool, gffcompare, part of the well known suite of isoform assembly and analysis tools Cufflinks. gffcompare at one time supported the notion of a “fuzz” parameter wherein intron boundaries with isoforms were allowed to be off by a certain length. This mode was disabled in more recent versions. To our knowledge no other tool does this. We re-enabled this mode specifically for this paper’s work. This allowed us to apply the fuzz approach we took with individual splice junctions to the isoform level. Specifically, we focused on comparing intron-chains between two isoforms. That is we ignored the start/end exons and restricted the comparison to only the coordinates of the introns.

In addition, we added a parameter which forces gffcompare to load its “reference” and “query” sets of isoforms in exactly the same way, applying the same deduplication approach to both. This ensured that the same pair of samples run in one order would be the same in the reverse. The sensitivity and precision numbers are derived from intron-chain comparison.

All comparisons are made via gffCompare (updated version of cuffCompare from Cufflinks) and are exact matches.

Union of Annotations is made up of: Gencode V29, Gencode V26 (GTEx), RefSeq HG38 (as of early 2019), FANTOM-CAT 6 (lncRNA + Gencode).

**Supplemental Note 3. Training simulation dataset pipeline**

Each of the four datasets (Oxford FL, Oxford non-FL, PacBio FL, PacBio non-FL) were simulated by taking the error profile generated by SURVIVOR and using SURVIVOR's "simreads" command from the original Minimap2 alignments and using this with the set of transcript sequences from Gencode V28 to produce synthetic reads which were then aligned back to the genome using Minimap2. This process was run five separate times. Per-simulated run error categories are defined in the main text as A-E at the start of the Simulation section.

1. Original NA12878 Oxford/PacBio long reads aligned against Gencode V28 transcript sequences
2. SURVIVOR "scanreads" extracts technology specific error profile from alignments in 1. Using a read length cutoff of >=100
3. SURVIVOR "simreads" then simulates new reads based on steps 1. & 2.
4. Synthetic long reads from step 3. are then mapped back to HG38 using Minimap2 using the same parameters across all runs

Steps 3-4 were run 5 times per technology for both full-length and fragments producing a total of 20 runs. Any transcript that was consistently novel across all simulation runs was assigned to the novel class. Any transcript that was consistently in all of the error classes, but not consistently novel, was assigned to the recurrent-error class. Any transcript that was not in either the consistently novel or recurrent error classes was but had been in at least one or more error categories in one or more of the runs was assigned into the non-recurrent error class. Finally any transcript not previously filtered for was assigned a problem-free category.

The flow of decision for categorizing a single simulated transcript is as follows:
1. Consistently novel across all simulations runs? Novel (end)
2. Not in Novel and consistently in all 3 error categories across all simulation runs? Recurrent-error (end)
3. Not in Novel/Recurrent-error but was categorized in >= 1 error category in >= 1 simulation run? Any-error (end)
4. Must be Problem-free (end)

This logic is defined in the `compare_matching.sh` script in the associated Github repository.

The original alignment of the Oxford NA12878 sample was using this command line:

```
minimap2 -ax splice -uf -k14 -t 16
./referenceFastaFiles/dna/GRCh38_full_analysis_set_plus_decoy_hla.fa
./NA12878-DirectRNA.pass.dedup.NoU.fastq
```

All simulations were run with the same parameters and the same reference.

**Supplemental Note 4. Counting results of predictions on NA12878**

Once models were trained for the four cases, these were used to predict the labels of the original Minimap2 alignments of the same samples. The RandomForest probabilities for each class for each alignment were then tabulated and a match was recorded if the class with the highest probability matched the training label or the probability the matching class was within 0.02 of the class with the highest probability.

**Supplemental Note 5. NA12878 & SKBR Custom Tracks in the UCSC Genome Browser**

NA12878 Oxford:
http://snaptron.cs.jhu.edu/data/longtron/oxford.NA12878-DirectRNA.bam.bed.sorted.bb

NA12878 PacBio:
http://snaptron.cs.jhu.edu/data/longtron/pacbo.SRR1163655.sorted.bam.noHLA.bed.bb

SKBR3 PacBio:
http://snaptron.cs.jhu.edu/data/longtron/SRR7346977_subreads.bam.nohla.bed.bb

**Supplemental Note 6. Features used in the Random Forest training/prediction**

Feature Files:

- Repeat Masker overlap: hg38_repeatmasker_rmsk
- Tandem Repeat Finder (TRF) overlap (subset of 1)): simple_repeats_hg38
- Splice motif count: hg38_splice_motifs.all.bed.bgz
- Exon/intron statistics: gencode.v28.basic.annotation.exons.stats.bed (locus stats), gencode.v28.basic.annotation.exons.perbase.counts.bgz
- # of overlapping reads/transcripts: gencode.v28.basic.annotation.transcripts.perbase.counts.bgz
- # of common 150 SNPs overlapping: snp150Common.combined.sorted.bed.no_bad_chrs
- GC content: gc5Base.bg.clean
- General Mappability: k24.Umap.MultiTrackMappability.sorted.bg
- Local Mappability: gv28.local_mappability.coords.bed
- Segmental Duplications: segmental_dups_hg38.sorted

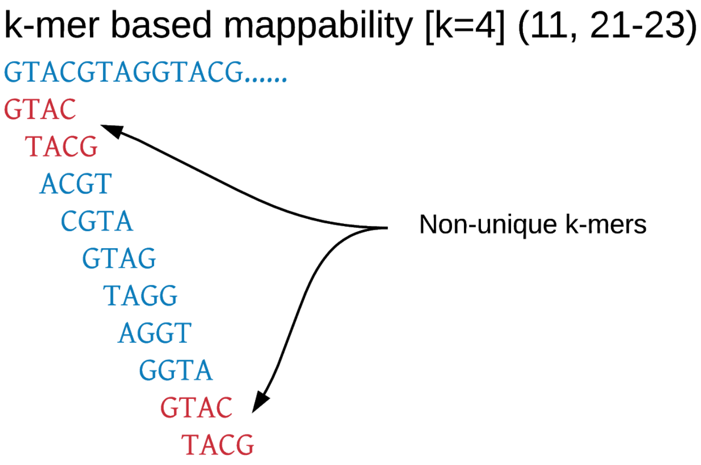


**Figure S1.** Kmer mappability. Mappability is based on k-mers, k=24 for umap multi-tracking mappings and k=10 for local region mappings. This is for features used in the random forest: 11, and 21-23.

**Figures S2.** ROC plots of random forest accuracy with features

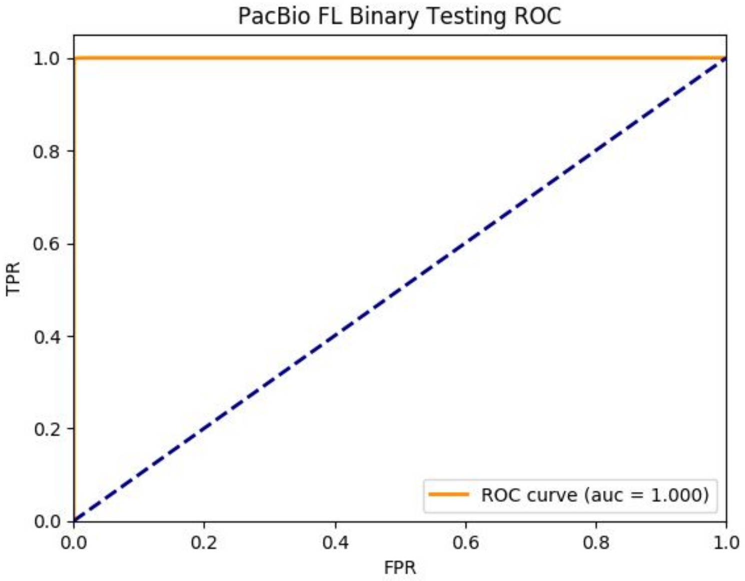


**Figure S2-A.** Oxford FL Binary Class ROC on Testing (held-out) data

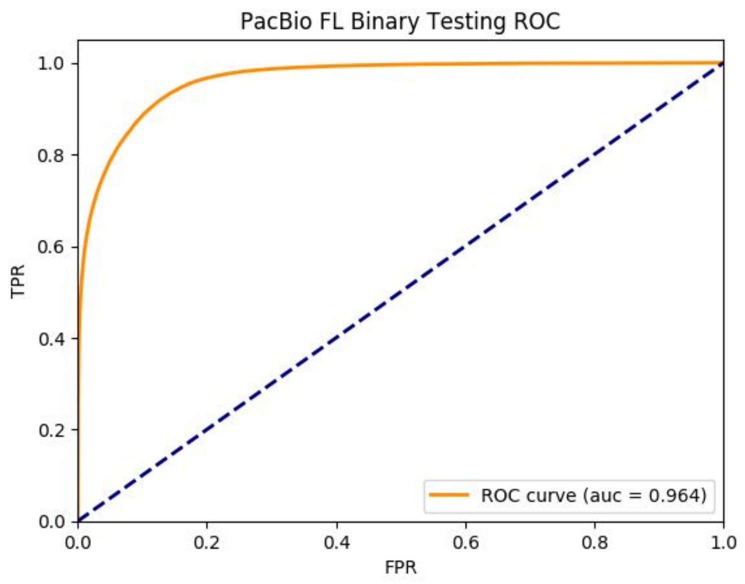


**Figure S2-B.** Oxford non-FL Binary Class ROC on Testing (held-out) data

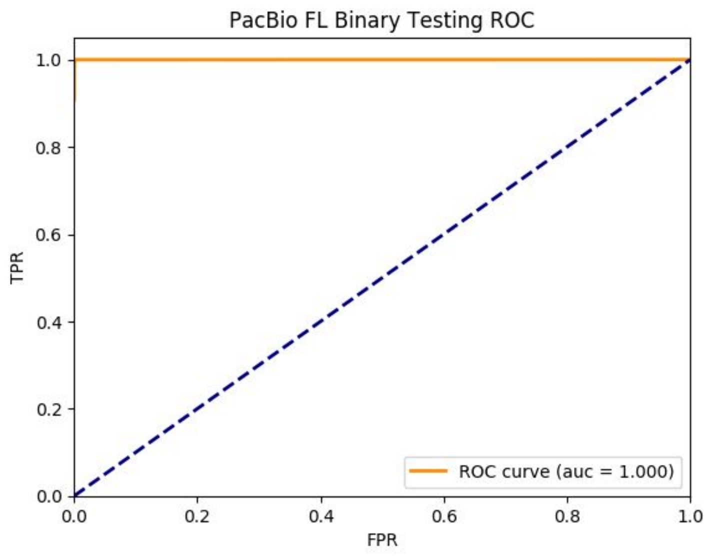


**Figure S2-C.** PacBio FL Binary Class ROC on Testing (held-out) data

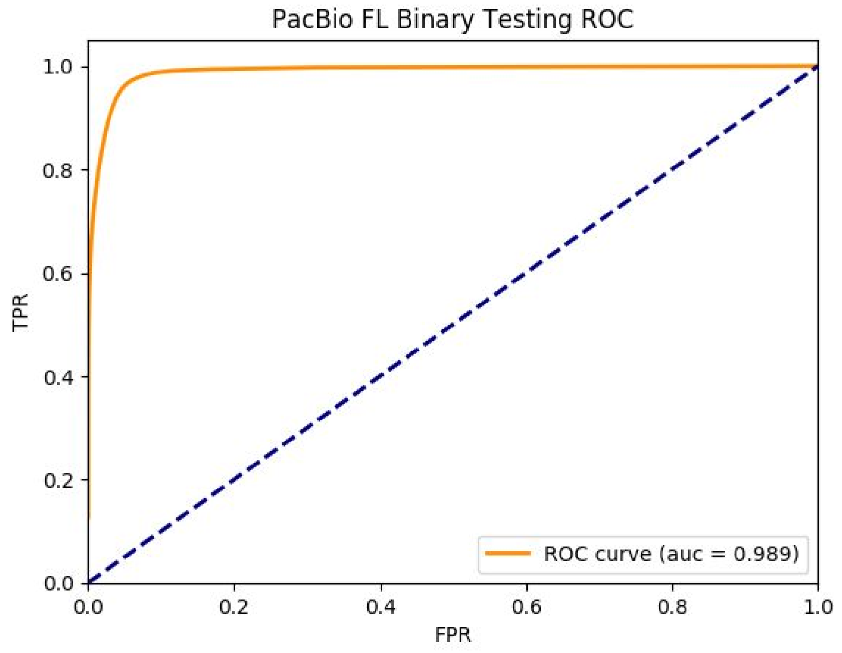


**Figure S2-D.** PacBio Non-FL Binary Class ROC on Testing (held-out) data

**Table S3.** NA12878 Alignment Class Recall

| Dataset | Total Reads Categorized | Problem-free | Any error | Recurrent error | Novel |
|---|---|---|---|---|---|
| **Oxford full length** | 2,749,480 / 5,284,512 | 89% (2282118 / 2552987) | 17% (453539 / 2646694) | 18% (12421 / 70490) | 10% (1402 / 14341) |
| **Oxford fragment** | 2,406,066 / 5,492,870 | 44% (1519189 / 3468504) | 62% (883498 / 1418051) | 1% (3221 / 292757) | 0% (158 / 313558) |
| **PacBio full length** | 228,317 / 261,944 | 98% (224685 / 228633) | 12% (3493 / 29760) | 3% (72 / 2072) | 5% (67 / 1479) |
| **PacBio fragment** | 120,756 / 285,799 | 48% (115026 / 240716) | 21% (5399 / 26276) | 38% (238 / 630) | 1% (93 / 18177) |

**Table S4.** NA12878 Alignment Class Precision

| Dataset | Total Reads Categorized | Problem-free | Any error | Recurrent error | Novel |
|---|---|---|---|---|---|
| **Oxford full length** | 2,749,480 / 5,284,512 | 51% (2282118 / 4459215) | 67% (453539 / 675162) | 34% (12421 / 36293) | 1% (1402 / 113842) |
| **Oxford fragment** | 2,406,066 / 5,492,870 | 62% (1519189 / 2434882) | 29% (883498 / 3043506) | 23% (3221 / 13835) | 24% (158 / 647) |
| **PacBio full length** | 228,317 / 261,944 | 88% (224685 / 254032) | 51% (3493 / 6835) | 42% (72 / 171) | 7% (67 / 906) |
| **PacBio fragment** | 120,756 / 285,799 | 83% (115026 / 137759) | 11% (5399 / 48902) | 0% (238 / 97771) | 7% (93 / 1367) |

Totals in the table are per-category and based on the total number of alignments that were predicted to have that class label. Totals are the same between precision and recall and are repeated for convenience.

**Table S5.** Intron Chains in Annotation [exact (fuzz) percent matching]

| Annotation | Total Intron Chains | Illumina NA12878 | Illumina SKBR3 | Oxford NA12878 (OX-RAW) | Oxford NA12878 FLAIR | PacBio NA12878 (PB-RAW) | PacBio NA12878 FLAIR | PacBio SKBR3 (PB-SKBR3) |
|---|---|---|---|---|---|---|---|---|
| **Gencode V29** | 199,381 | **17.0% (29.8%)** | 12.0% (22.7%) | **35.9% (44.1%)** | 15.2% (25.9%) | 17.5% (26.2%) | 7.1% (14.2%) | 27.2% (37.3%) |
| **Union of Annotations** | 1,098,511 | **15.4% (26.3%)** | 12.4% (20.7%) | **36.2% (44.2%)** | 13.0% (22.0%) | 19.0% **(26.2%)** | 7.6% (12.7%) | 28.1% (38.3%) |

**Table S6.** Improvement of intron-chain matches from problem free predictions

| **Dataset** | **Intersection** of FL & Fragment Problem Free Predictions vs. **Union of Annotation** | **Union** of FL & Fragment Problem Free Predictions vs. **Union of Annotation** |
|---|---|---|
| **NA12878 Oxford vs. Annotation** | 82.7% (97.5%) | 82.8% (96.9%) |
| **NA12878 Pacbio vs. Annotation** | 57.2% (86.7%) | 46.3% (82.0%) |
| **SKBR3 Pacbio vs. Annotation** | 31.3% (67.3%) | 28.3% (64.6%) |
| **Annotation vs. NA12878 Oxford** | **19.9% (25.9%)** | **33.2% (40.7%)** |
| **Annotation vs. NA12878 Pacbio** | 13.7% (20.5%) | **18.8% (26.0%)** |
| **Annotation vs. SKBR3 Pacbio** | 22.4% (33.7%) | 27.9% (38.1%) |
